# A phenotypically plastic magic trait promoting reproductive isolation in sticklebacks?

**DOI:** 10.1101/587675

**Authors:** Monica V. Garduno Paz, Felicity A. Huntingford, Sean Garrett, Colin E. Adams

## Abstract

This study identifies one possible mechanism whereby gene flow is interrupted in populations undergoing evolutionary divergence in sympatry; this is an important issue in evolutionary biology that remains poorly understood. Variation in trophic morphology was induced in three-spined stickleback by exposing them from an early age either to large benthic or to small pelagic prey. At sexual maturity, females given a choice between two breeding males, showed positive assortative mate choice for males raised on the same diet as themselves. The data indicate that this was mediated through a preference for males with trophic morphology similar to that of fish with which the females were familiar (from their pre-testing holding tanks). In trials where the female did not choose the most familiar male, the evidence suggests that either she had difficulty discriminating between two similar males or was positively choosing males with more extreme morphologies (more benthic-like or pelagic -like). This study has shown for the first time that expression of a plastic trait induced at an early age, not only results in specialisation for local foraging regimes but can also play a significant role in mate choice. This is equivalent to an environmentally induced, plastic version of the “magic traits” that promote ecologically-driven divergence in sympatry, hence the proposed descriptor “plastic magic trait”.

## Introduction

The process whereby gene flow is interrupted in populations undergoing evolutionary divergence when in sympatry is an important issue in evolutionary biology that remains poorly understood. One potential mechanism is where an ecologically important trait which is under divergent selection also contributes to reproductive isolation and is thus a so-called “magic trait” (Gavrilets 2004; Servedio et al. 2011). Although evidence of such traits in nature is sparse (Servedio et al. 2011), magic traits are usually envisaged as inherited and linked to mate choice through pleiotropy. However, much research interest has focussed on the possible role of phenotypic plasticity in the initiation of evolutionary change through the development of discrete alternative phenotypes (West-Eberhard 1989, 2003; Fitzpatrick 2012; Skulason et al. 2019). If expressed alternative phenotypic traits induced by the environment through plasticity, also form part of the mate choice system of the diverging organism, then assortative mating resulting from mate choice based on such traits have the potential to generate reproductive barriers between individuals expressing different phenotypes (Fitzpatrick 2012). Such traits may thus act as a magic trait without the requirement of pleiotropy.

Discrete alternative phenotypes associated with foraging, or trophic polymorphisms (*sensu* Skúlason & Smith (Skulason and wund 1995), have been strongly implicated in sympatric speciation events (Dieckmann and Doebeli 1999). Divergent morphological traits are often the result of foraging conditions experienced during development (Day and McPhail 1996; Adams et al. 2003), so can only result in evolutionary change if mechanisms exist that result in gene pool segregation (West-Eberhard 1989; Smith and Skulason 1996; Schluter 2003). Here we explore one possible mechanism, namely morph-specific mate choice by breeding females. Using the three-spined stickleback (*Gasterosteus aculeatus*) as a model system, we present an example of a developmentally-plastic, trophic specialisation acting as a magic trait generating reproductive isolation and suggest a mechanism through which this comes about.

Trophic polymorphism is particularly common among freshwater fishes including sticklebacks; it often takes the form of co-existing but discrete phenotypes with morphological and behavioural specialisations for feeding on benthic invertebrates in the littoral zone or zooplankton in the pelagic zone (Skulason and Smith 1996; Adams and Huntingford 2002a; Proulx and Magnan 2004). Typically, the benthic form is robust, with a large mouth, small eyes and few short gill rakers, while the pelagic form is lightly built, with a relatively small mouth, large eyes and longer and more numerous gill rakers (Adams et al. 1998; Adams and Huntingford 2002a). Although in some cases the two sympatric forms are fully reproductively isolated, more commonly reproductive isolation is partial, weak or non-existent (Schluter and McPhail 1992; Hendry et al. 2009). Sticklebacks also exhibit scope for the expression of characteristics under phenotypic plasticity (Wund et al. 2008, 2012; Garduño-Paz et al. 2010; Baker et al. 2013) including plasticity in morphological characteristics that define sympatric forms described from the wild (Day and McPhail 1996; Garduño-Paz et al. 2010).

The main aim of this study was, having induced variable trophic morphology in three-spined sticklebacks from a single population by manipulating early feeding regimes, to determine whether these plastic, diet-induced differences in trophic morphology were associated with different patterns of mate choice. A second aim was to seek possible behavioural mechanisms that might explain the observed patterns of mating.

## Methods

### Diet Treatments

240 juvenile three-spined sticklebacks fry (5-9 mm length) were collected by dip nets from a small freshwater pond in Scotland (56 ° 3’N; 004°21’W) and transported to rearing facilities at the Scottish Centre for Ecology and the Natural Environment (SCENE), Glasgow University, Loch Lomond. Fish were assigned randomly in groups of 40 to 6 rearing aquaria (21L) and raised in the laboratory for 11 months, during which time they were fed twice daily to satiation on one of two diet treatments known to induce differences in trophic morphology (Day and McPhail 1996). Half of the groups were fed on frozen *Daphnia* spp in a bag hanging at the water surface, simulating pelagic prey; the rest were fed on frozen chironomid prey placed on the bottom of the tank, simulating benthic prey.

### Analysis of Induced Morphological Differences

After 10 months, the sticklebacks were anaesthetised with benzocaine and photographed on their left side with a Canon EOS digital 350D camera (8.0 megapixels). Female fish that were gravid and male fish which were coloured were identified as sexually mature. All fish used in mate choice experiments were re-photographed at 11 months immediately following the mate choice experiments. Body shape was quantified on the basis of 20 landmarks (Figure S1), placed using the program “tpsUtil” ad digitised using the program “tpsDIg2” (Rohlf, 2006). Generalized least squares procrustes superimposition was used to translate, scale and rotate raw landmark coordinates to minimize the summed, squared, inter-landmark distances among fish; this procedure removes the effect of fish size (Rohlf & Slice, 1990). Relative Warp Analysis was conducted on the partial warp scores to reduce the number of informative shape variables (Rohlf, 2006). The second relative warp (analogous to a Principal Component), which explained 13% of the total shape variation, separated traits typical of pelagic and benthic feeders (Day and McPhail 1996; Figure S1).

### Mate Choice Trials

Twenty eight females (21 from the chironomid diet and 7 from the *Daphnia.* diet) and 36 males (21 from the chironomid diet and 15 from the *Daphnia* diet) were used in trials of mate choice. Female mate choice was examined using a well-tested methodology, widely used in previous studies (Milinski and Bakker 1990; Kraak and Bakker 1998; Boulcott et al. 2005; Rick et al. 2006; Rick and Bakker 2008; Heuschele et al. 2009) in which a single gravid female was placed alone in an aquarium (35 × 25 × 20 cms, screened on 3 sides), allowed to settle for 12h and was then presented simultaneously with two breeding males in equally-sized sections (25 × 35 × 20 cms) of an adjacent aquarium and thus not subject to olfactory cues. During trials, females could see both males, but the females had no olfactory contact with males and the two males were separated by an opaque partition and so did not have visual contact with each other. In any trial, the female was presented with one *Daphnia*-fed and one chironomid-fed male. To avoid effects of size and familiarity with specific males, the two males in any given trial were size-matched as far as possible and importantly taken from a different rearing tank from the female. To avoid females potentially making choices based on nest construction, males were not provided with nesting material.

Each trial lasted for 5 minutes, during which, the time the female spent on the side of the tank adjacent to each male was recorded. Three replicates of each pairing trial were conducted, swapping the male position each time. A female was deemed to have chosen a male if she spent at least 60% of the total time of the trial near that male. Association time has been shown to be a strong predictor of eventual mate choice in this species in a number of other studies (McLellan and McPhail; Rowland et al. 1995; Milinski et al. 2005; Rick and Bakker 2008). Males and females were used maximally in four trials on different days; males were re-used in fresh combinations so that the female was never exposed to the same pair of males. Although male pairs were matched in size (by fork length) as nearly as possible, small discrepancies between pairs remained. Retrospective analysis detected no significant difference in body size (fork length) between chosen and rejected males (mean ± SE size differences between accepted and rejected males =0.04 cm ± 0.02 paired t test: t = 1.68, p = 0.10).

Body shape, as defined by the second relative warp varied markedly both between and within diets (Fig.1). Effects of sex (ANOVA: F_1,60_=3.11; p=0.08) and of sex by diet (ANOVA: F_1,60_=2.86; p= 0.09) were not significant. However, there was a highly significant difference in morphology between the chironomid fed (mean ± SE = 6.753 ± 0.23) and *Daphnia* fed sticklebacks (mean ± SE = 4.82 + 0.34. ANOVA: F_1,60_= 22.2; p< 0.0001). The higher scores of fish fed on the benthic diet reflected shorter heads, shorter maxillary bones, smaller eyes and deeper bodies. This score was transformed to create only positive values (by adding 6 and multiplying by 100) and hereafter this dimension of shape variation is referred to as the pelagic-benthic (PB) shape score. Lower PB scores indicate shapes tending towards a more typical of a pelagic foraging fish; higher scores tending towards a more benthic foraging fish shape (Fig.1). To enable testing of the body shape of chosen and rejected males, the difference in PB shape score between the chosen male and the rejected male for each pair was determined as: male PB score difference =PB score chosen male – PB score rejected male. Thus a large positive number in this metric indicates the choice of a male from the pair with a higher PB score; thus tending towards a shape more typical of a benthic foraging fish. Conversely a large negative number in the difference between chosen and rejected male PB scores indicates a choice of male with a shape tending towards that typical of a more pelagic forging fish.

**Figure 1.**
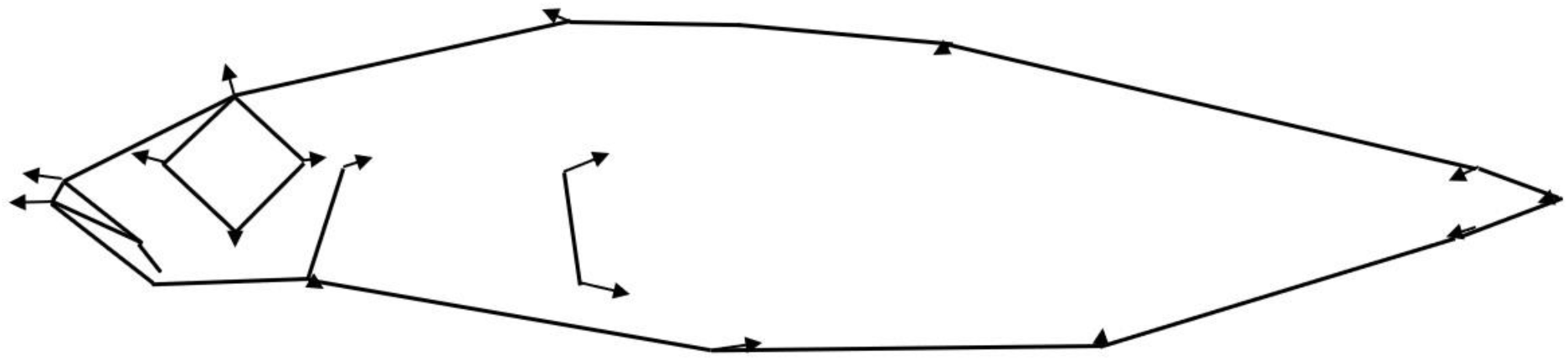
The landmark configurations used in the morphometric analysis of trophic morphology in sticklebacks. The landmarks are connected to aid visualisation of fish shape. Arrows represent vectors describing deformations that change the mean shape of sticklebacks fed on benthic prey compared with the mean shape of those fed on pelagic prey.

In nature, female sticklebacks review a number of males before selecting a nest in which to lay her eggs and the females in this study were always tested with different pairs of unfamiliar males. However female identity was included as a random factor in mixed effects models examining female mate choice. A total of 96 trials of mate choice by females was analysed using “R” (R Development Core Team 2017).

## Results

### Mate Choice in Females Exposed to Benthic and Pelagic Diets

To test for the effect of the previous diet experience of the female on female mate choice, the PB score difference between male pairs in each trial was categorised as either a negative value, indicating the choice of a male with a more pelagic-like body shape, or positive, indicating a choice of the male from the pair with a more benthic-like body shape. The previous diet experience of the female (benthic (chironomid) or pelagic (*Daphnia*) diet exposure) was used as a fixed factor and female I.D. as random factor, in a mixed effects model with a binomial (positive or negative male PB score difference) probability distribution. Female diet exposure significantly predicted the choice of male body shape in pairwise tests (p<0.01) indicating non-random mating on the basis of previous diet exposure. However the total explained variance was low (6.8%) and this effect was mostly driven by females with previous exposure of a pelagic prey diet choosing the male with the more pelagic body shape (a negative male PB score difference) on 66% of occasions (Table 1). In contrast, females exposed to the benthic prey diet chose the male from the pair with the more benthic body shape (a positive male PB score difference) on only 54% of occasions.

**Table 1.** The frequency with which the male with the higher pelagic-benthic score (more benthic-like) was chosen and rejected by female sticklebacks exposed to alternative diets (a pelagic like diet (*Daphnia*) and a benthic-like diet (chironomid larvae).

| Female diet | Male with more positive P-B score | | $\chi^2$ DF P |
| --- | --- | --- | --- |
|  | Chosen | Rejected |  |
| Benthic | 34 | 29 | 0.4; 1; >0.53 |
| Pelagic | 9 | 24 | 6.8; 1; <0.01 |

### Behavioural Mechanisms Of Mate Choice

To explore possible behavioural mechanisms for the observed female preference by diet, mate choice data were analysed in more detail. To test the possibility that females are making a choice of male based on their own body shape, the male PB score difference in each trial, categorised as either a negative value, (indicating the choice of a male of a more pelagic-like body shape), or a positive value (indicating a choice of the male with a more benthic-like body shape) was modelled using female PB score as a fixed factor and female I.D. as random factor, in a mixed effects model with a binomial probability distribution. Female body shape did not predict the choice of male (*p*<0.45). Thus the assortative mating effect predicted by female previous diet exposure does not appear to be driven by the body shape of the female.

Another possible behavioural mechanism by which rearing diet might influence a female stickleback’s mate choice is through previous experience of the fish with which she was reared, whose shape will, on average, reflect their common rearing diet. To test this possibility, we took advantage of the variability in PB scores between rearing tanks on a given diet. The male PB score difference in each trial, categorised as either a negative value, or a positive value was modelled using the mean PB score of fish in the tank from which the female was drawn, as a fixed covariate and female I.D. as random factor, in a mixed effects model with a binomial probability distribution.

The mean PB score of fish from the tank from which the female originated significantly predicted the female’s choice of male (*p*<0.02; r^2^= 6.9%). Indicating that females are choosing of males on the basis of body shapes with which they are familiar. Despite this a considerable amount of the variation in the choice of male was not explained by the body shape with which they were familiar.

One possible explanation is that in trails where females chose males that were not closer in body shape to those with which she was familiar (from the same rearing tank) and thus she may have had difficulty discriminating between body shapes. If this was the case, then one expectation would be that in such trials the two males are likely to be closer in body shape (similar PB scores) to each other than in trials where the female chose the male closest to that with which she is familiar. This was tested; the PB score difference between males in trials where the female chose the most familiar shape (2.22 ± 1.36; mean ± S.D.) was significantly greater (t test: *p=*0.041) than the PB score difference between males from trails where she chose the less familiar male (1.85 ± 0.94). This indicates that in at least some trails females may have had difficulty distinguishing between males of similar body shape.

Another possible explanation for the outcome of those trails where the female did not chose the male with a body shape closest to the mean of fish that she was familiar with, is that she may avoid choosing males with extreme body shapes even if they are closer to the body shape with which she is familiar. To test this the Extreme Shape Index was calculated as the deviation in body shape for chosen and rejected male from the average of all fish combined (that is the PB score of each fish - the mean of all fish combined irrespective of sign. As these data deviated significantly from normality, the Extreme Shape Index for chosen and rejected males from trials where the female chose the male with a shape that she was less familiar with, were compared in a paired Wilcoxon test. In these trials the chosen male was much more likely to have a more extreme body shape (i.e a higher Extreme Shape Index) (2.09 ± 1.08; mean ± S.D.) than the rejected male (1.01 ± 0.86).

## Discussion

Our results confirm the findings of previous studies demonstrating a plastic response of morphological traits to rearing diet in three-spined sticklebacks (Day and McPhail 1996). More significantly, they have demonstrated for the first time that exposure to different diets during the juvenile phase can influence the mating preferences shown by breeding females. Thus, females reared on the pelagic diet tended to prefer the male with a more pelagic-like morphology; females reared on a benthic diet however mated randomly with respect to trophic morphology. Thus there is partial assortative mating by diet-induced phenotype. Unlike the case of assortative mating on the basis of diet specialisation in the mustard leaf beetle, which appears to use olfactory cues to identify mates (Geiselhardt et al. 2012) the sticklebacks in this experiment only had visual cues are available to them. However, it is quite possible that olfactory cues might also have affected mate choice had they been available. In addition, the effect reported here did not result from female familiarity with specific individual males, as females were never tested with males from the same rearing tank.

Additionally we show that mate choice was not dependent directly of the female’s own trophic morphology. Arguably, this is not surprising, since it is difficult to see how a female stickleback could know what her own morphology is like. Instead the differences in mate choice must be a consequence (direct or indirect) of the experience of being raised on a pelagic or a benthic diet. Making use of the significant variation in morphology between and within rearing tanks exposed to different and the same diets, we show that the expressed morphology of other fish with which the female is familiar (from the same rearing tank) is a good predictor of mate choice. It is highly likely that in the wild also sticklebacks grow up with fish exploiting a similar diet to themselves and thus with similar diet-induced morphology, as individuals exploiting the same foraging resources are more likely to come into contact with each other, than those that do not share a common diet (Garduño-Paz and Adams 2010).

Despite a clear tendency for assortative mating by trophic morphology, females quite often made the opposite choice. This was most often the case when the difference between the two males was relatively small, but also occurred when if the morphology of the predicted choice male was of an extreme benthic or pelagic-type morphology. One can envisage at least two plausible mechanistic explanations for this, which are not mutually exclusive. It may be that, rather than responding to familiarity *per se*, females have learned about the foraging efficacy of fish with the range of morphologies that she has experienced during development. If this were the case, then females might actively choose males of a more extreme morphology, which may well be more efficient at foraging on the two alternative diets presented, even if this morphology is less common in her previous experience familiar to her. We are not able to test directly this possibility using the data from this study.

Although coexisting trophic morphs are thought to be an important step in evolutionary divergence in sympatry (Skulason et al. 1999), speciation however is unlikely to be completed without some mechanism for morph-specific assortative mating (Skulason et al. 1999). Several routes though which this might occur have been suggested. For example specialist morphs might occupy different habitats. Olafsdottir and co-workers (Olafsdóttir et al. 2006) for example, showed that sticklebacks specialising in living in habitats with little vegetation had reduced nest building behaviour and as a result weed-living specialists from the same lake mated assortatively with other weed-living specialists when using nest quality as a mate choice criterion. Disruptive sexual selection is also known to play a significant role in the divergence of recently evolved African cichlid species (Stelkens et al. 2008). Here uniquely we demonstrate assortative mating on the basis of morphological traits that frequently express as discrete forms in the wild, have strong functional significance for resource acquisition (Adams and Huntingford 2002b) are thought to be under strong selection pressure and the expression of which is significantly modulated by plasticity effects. This result indicates that trophic morphology is both a plastic and a magic trait for sticklebacks, thus that pleiotropy may not always be required for traits to operate as magic traits.

## Acknowledgements

We thank Rona Brennan for technical support. M.V.G-P. was supported by a Mexican Council for Science and Technology (CONACYT) scholarship. The authors declare no conflicts of interest.

## Compliance with ethical standards

This study was conducted in accordance with UK legislation under Home Office Licence Number: PPL 70/8794.

